## Supplementary2 for "Fragmentation and Multithreading of Experience in the Default-Mode Network"

### Supplementary Results

#### Supplementary Table 1

**State Instructions**

| Block | Instruction |
|---|---|
| 1 | ..In this short movie, several different events will happen each with a different underlying context. Sometimes these contexts shift abruptly.. |
| 2 | ..Example (not related to the movie): Imagine a clip of person going into a restaurant to eat food. This would be a context and thus, a situation model of this context can have any events that normally occur within a context of a 'person eating in restaurant'. Even if you are seeing this from different angles or cuts, the central context is unchanged. Suppose, two crooks come into the restaurant and robs the cashier. This would mean the current context is changed, and hence your beliefs would be updated while watching this.. |
| 3 | ..This denotes an update to the current context, and you will update your understanding of what is currently happening in the movie..Similarly, many such updates will occur in this movie. Whenever you feel the current context is changing, respond by pressing the s key in your keyboard. That is, throughout the movie, until its over, whenever you feel there is a context change press the s key.. |

**Agent Instructions**

| Block | Instruction |
|---|---|
| 1 | ..In this short movie, different characters will be present. These characters will appear in a variety of contexts -alone or interacting with others. Throughout the movie, your initial beliefs about each character may change based on their actions, reactions, or dialogues.. |
| 2 | ..Example (not related to the movie): Imagine a clip showing a person who is initially portrayed as friendly and approachable. This first impression might lead you to believe that they are a kind and helpful individual. But, if a later scene shows this character refusing to help a friend in need, you might need to reassess your initial beliefs about this individual. This would be an update event where your beliefs about this character have changed.. |
| 3 | ..This denotes an update to a character, and you will update your understanding of what is currently happening in the movie..Similarly, many such updates will occur in this movie. Whenever you feel your belief about a character is changing, respond by pressing the s key in your keyboard. That is, throughout the movie, until its over, whenever you feel there is an update to your belief about the characters press the s key.. |

**Action Instructions**

| Block | Instruction |
| --- | --- |
| 1 | ..In this short movie, you will notice different actions occuring. As you watch the movie, you will develop some expectations about what is likely to happen based on these actions. There will be actions taking place in the movie that would make you update your expectations about the movie.. |
| 2 | ..Example (not related to the movie): Imagine a clip about a group of friends on a hiking trip. Initially, they're shown checking their gear and supplies, leading you to believe it's a well-planned trip. But, if a scene later shows an action resulting in all of them losing their backpacks in a river, your expectations for what will happen next could change drastically due to the action that occured. Maybe they'll face hardships, turn back, or have to find new supplies. This moment, when your expectations about the movie's course can change due to the actions involved, is what we're interested in.. |
| 3 | ..Such action events alters the possible actions that can occur in the movie, updating your belief about the course the movie is going to take and thus you will update your understanding of what is currently happening in the movie.. Whenever you feel your belief about an action is changing, respond by pressing the s key in your keyboard. That is, throughout the movie, until its over, whenever you feel there is an update to your belief about the actions press the s key.. |

Table S1. Instructions for belief updates for States, Agents and Actions, for the Movie participants. Instructions for Narrative had the same structure with terminology modified for listening rather viewing.

### Supplementary Fig 1

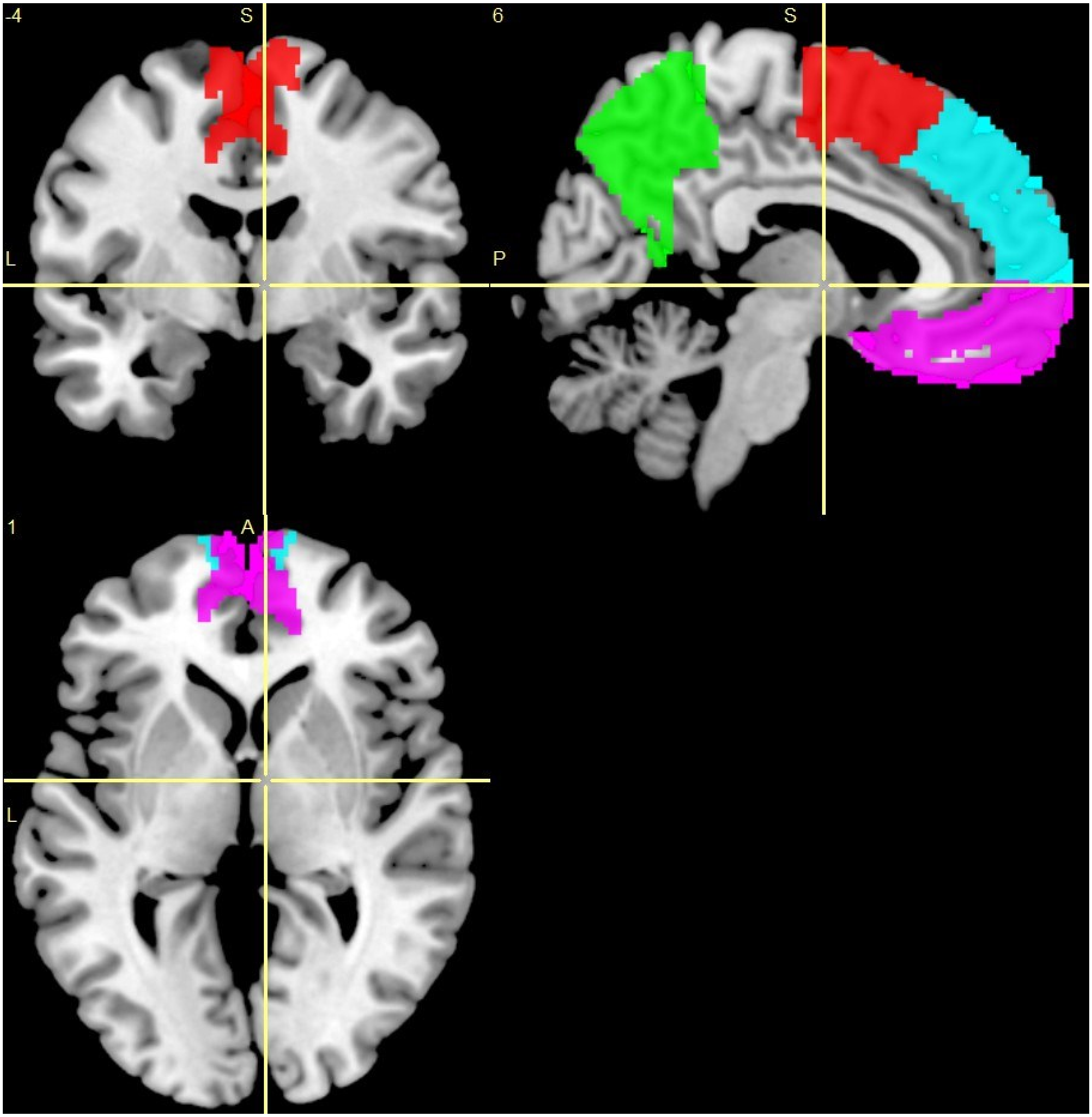


Supplementary Figure 1 : ROI Masks of Prefrontal regions and Precuneus.

#### Supplementary Fig 2

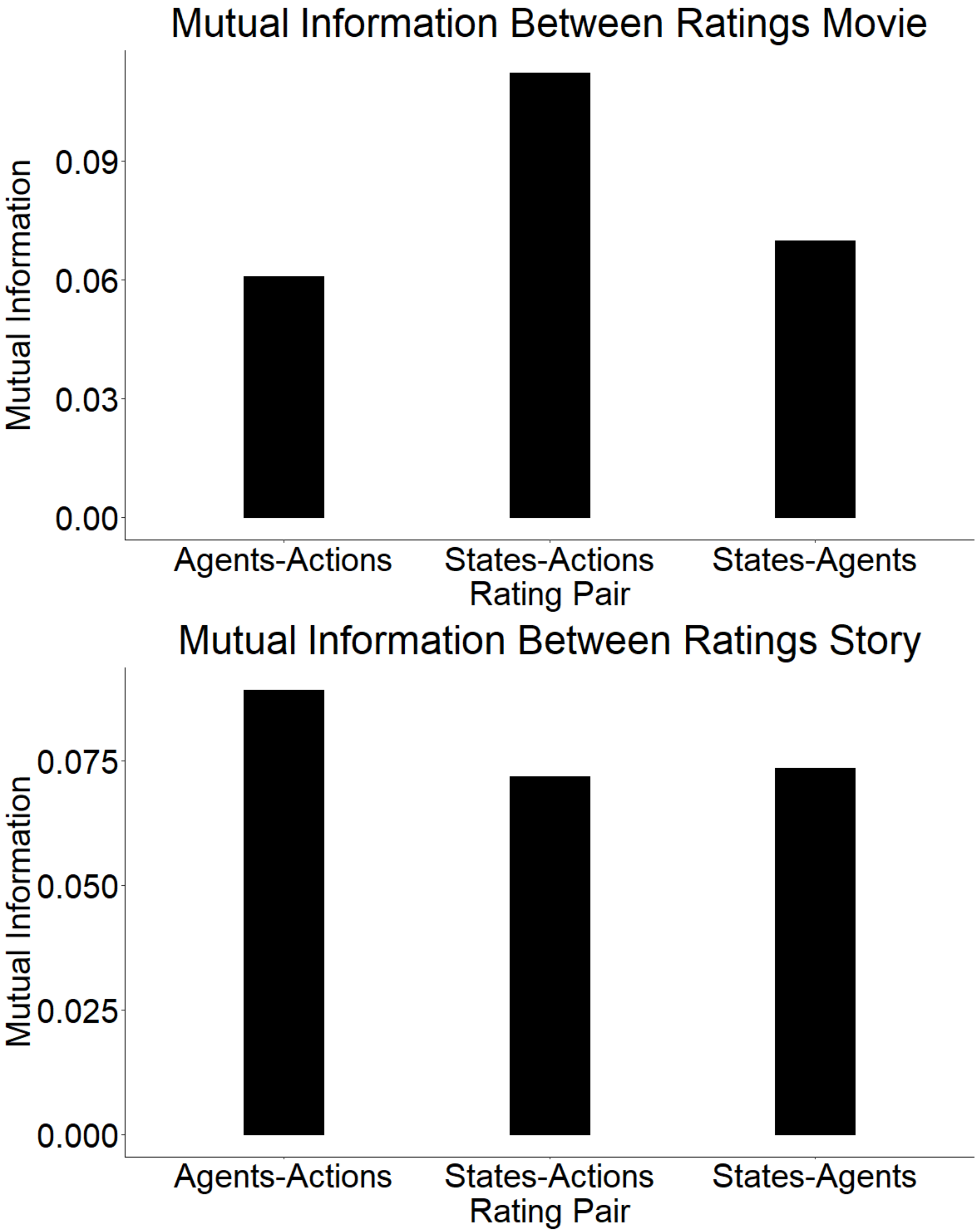


Supplementary Figure 2 : Mutual information between all three pairs of domain ratings in movie (Top) and story (Bottom). Values ranged between 0.05-0.09 (computed at bins of length $L^{\wedge 1/3}$, where L is the length of the time-series (ratings) used).

#### Supplementary Fig 3

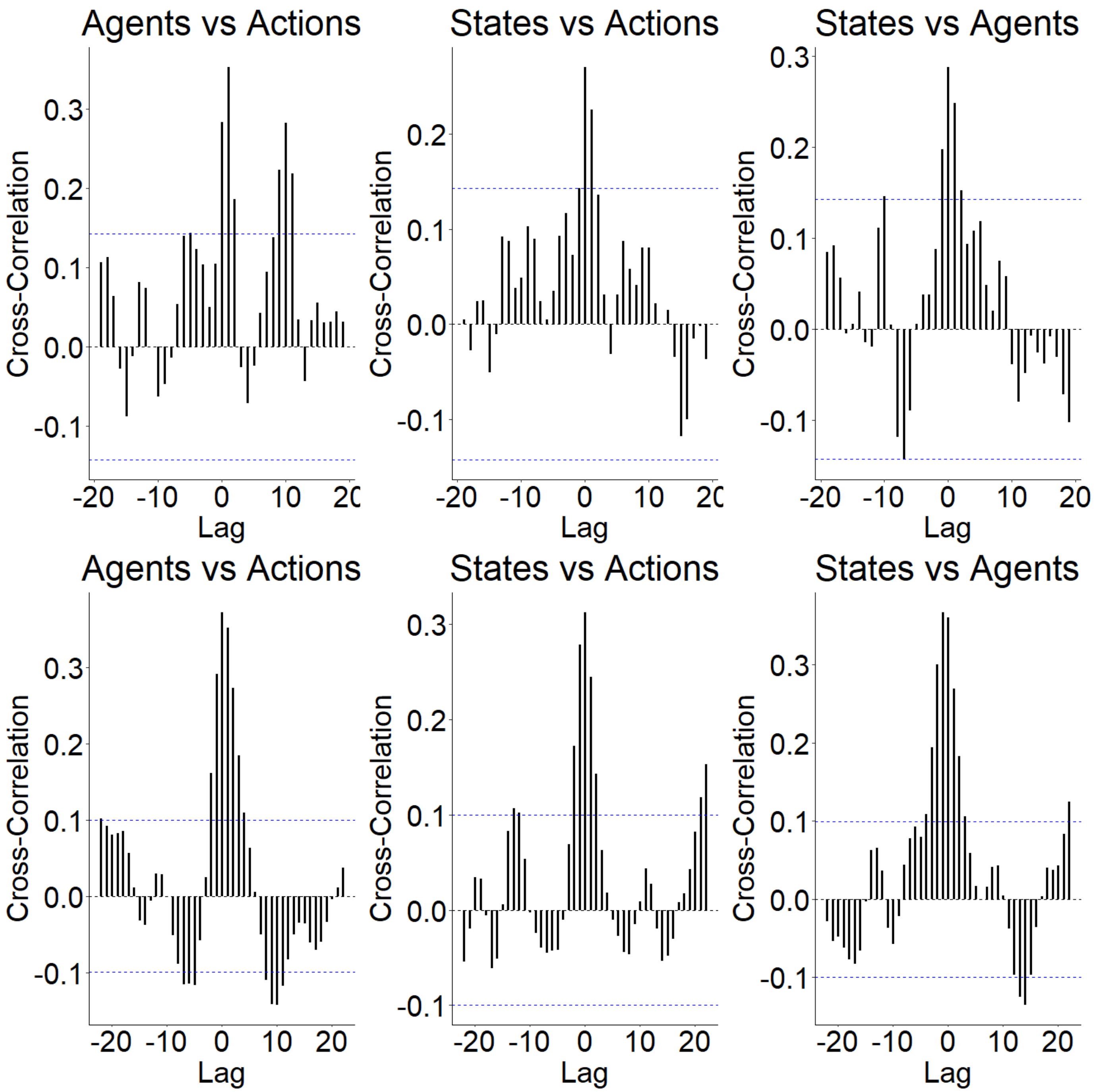


Supplementary Figure 3 : Cross-correlation between all three pairs of domain ratings in movie (Top) and story (Bottom). Blue dashed lines represents threshold for significant correlations.

### Supplementary Fig 4

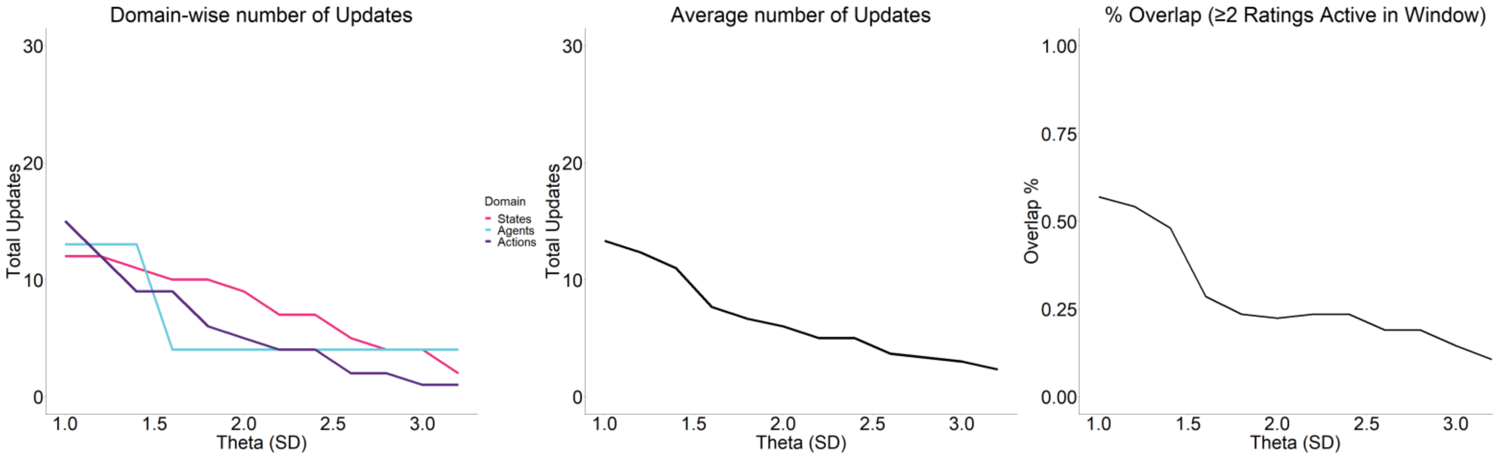


Supplementary  Figure 4 : Threshold Selection movie. **a**. We computed two separate metrics for the raw ratings (group-average), each reflecting two key aspects of an idealized rating ; number of available update points and overlap among the three domains. Any threshold, as it increases (e.g. here from 1 std to say 3.2 std) should increasingly lead to less and less updates surviving the stringent standard. This can be seen below for each domain in movie (left) as well as the averaged across domains (middle). But the number of updates alone does not dictate a selection criterion since more updates generally means, more overlap and thus makes the neural signal less specific to testing hypothesis of PFC sensitivity. For this we used an approach of constructing a window (10 TRs) and computing the number of overlaps. Overlaps here is taken as present in a window if >1 update occurs in the window. We slide this window throughout (i.e. all possible windows) and compute the fraction of windows with overlap to total windows (bottom)

### Supplementary Fig 5

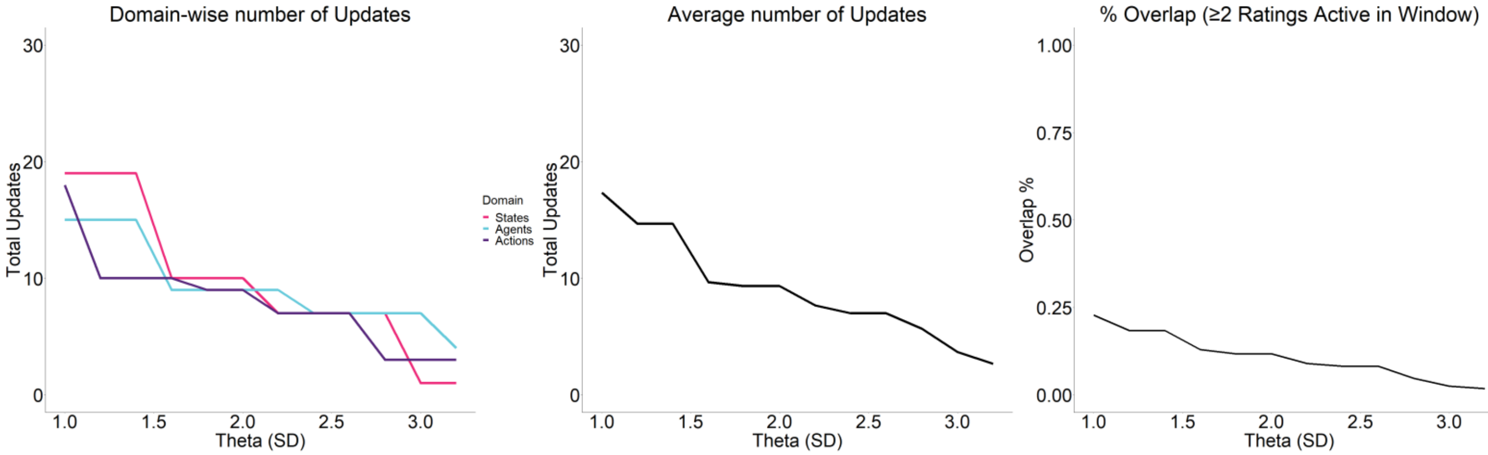


Supplementary  Figure 5 : Threshold Selection story. The exact same procedure of threshold selection used in movie (Supp Fig 4) was repeated in story showing **a.** total updates (left), averaged updates (middle) and overlap percentage (right) for various thresholds.

### Supplementary  Fig 6

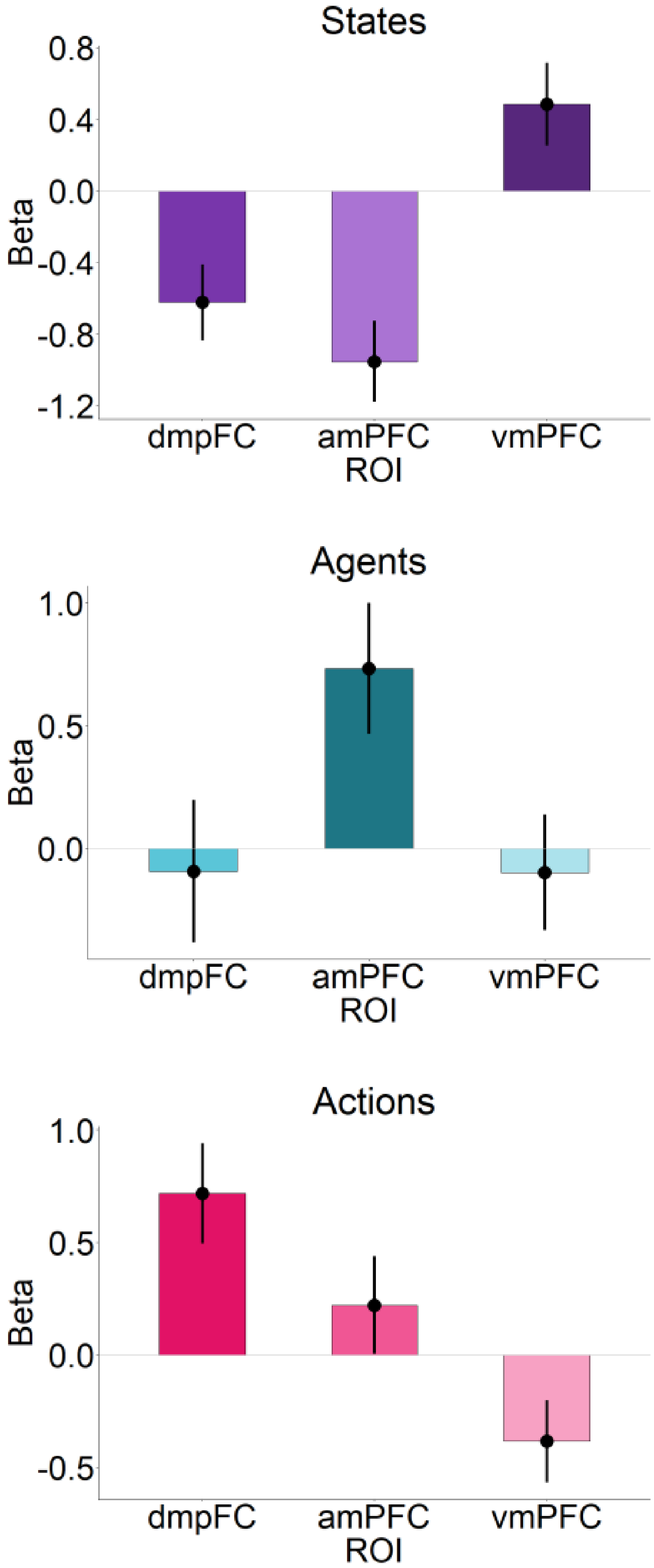
States
0.8
0.4
0.0
-0.4
-0.8
-1.2
Beta
dmpFC
amPFC
vmPFC
ROI
Agents
1.0
0.5
0.0
Beta
dmpFC
amPFC
vmPFC
ROI
Actions
1.0
0.5
0.0
-0.5
Beta
dmpFC
amPFC
vmPFC
ROI

Supplementary Fig 6 : ROI analysis confirming a domain-specificity within PFC regions. BOLD response to State updates was significantly higher in vmPFC than either amPFC or dmPFC (vmPFC > amPFC, t = 8.2548, p-value = 3.693e-13, d = 0.78, vmPFC>dmPFC, t = 5.1039, p-value = 1.404e-06, d = 0.48), while effects of Agent updates was highest in amPFC (amPFC > vmPFC, t = 4.0458, p-value = 9.718e-05, d =0.38, amPFC > dmPFC, t = 4.9594, p-value = 2.591e-06, d = 0.47) and Action updates highest in dmPFC (dmPFC > vmPFC, t = 5.0828, p-value = 1.536e-06, d =0.48, dmPFC > amPFC, t =3.5133, p-value = 0.0006434, d = 0.33). . Error bars show standard error of the mean.

### Supplementary Fig 7

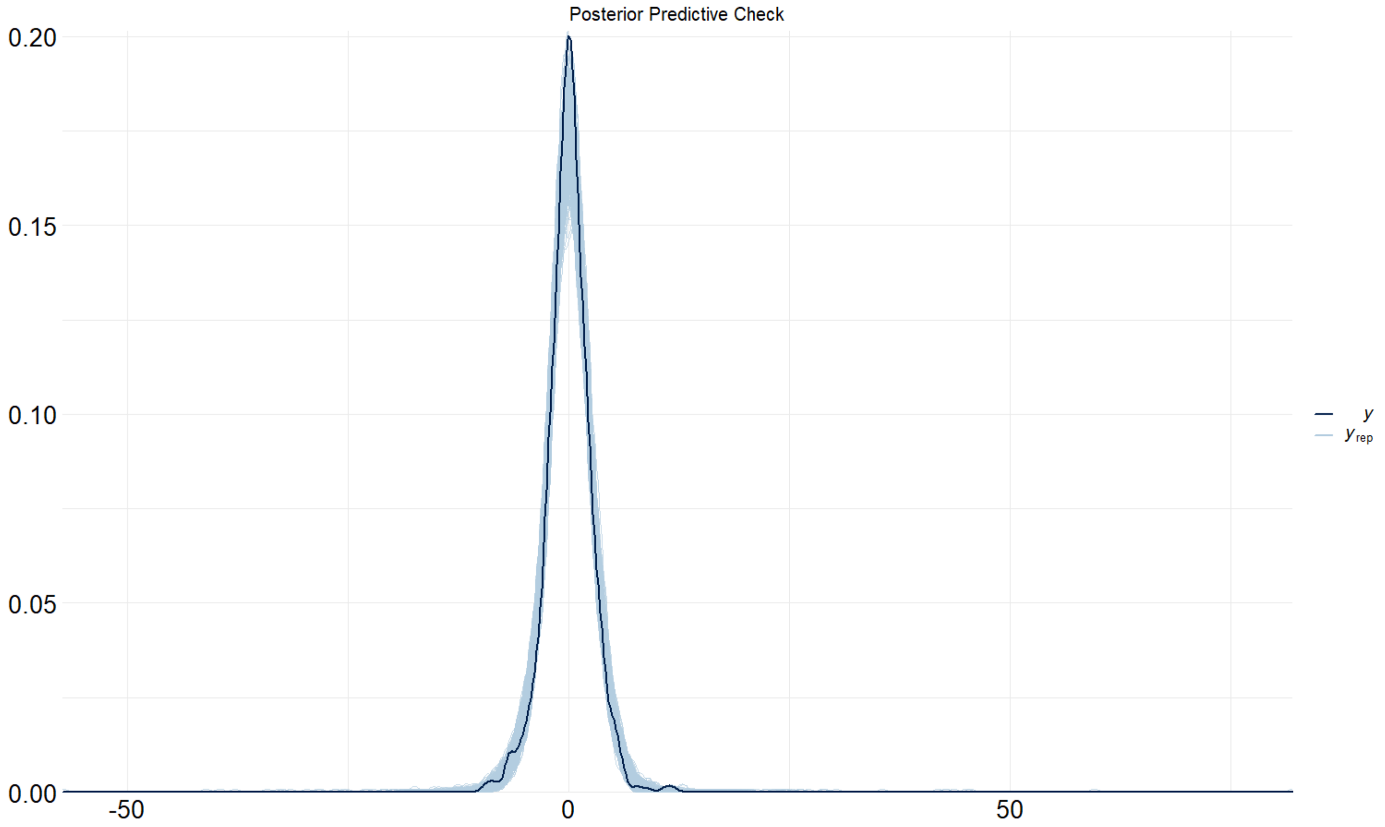


Supplementary Figure 7 : Posterior predictive draws (cyan) of 1000 simulations from the fitted model compared with data (black) for the information encapsulation Bayesian model for GLM ROI analysis (betas) within prefrontal regions during domain updates. The model had (Bayes adjusted) $R^2$ of 0.72.

### Supplementary Fig 8

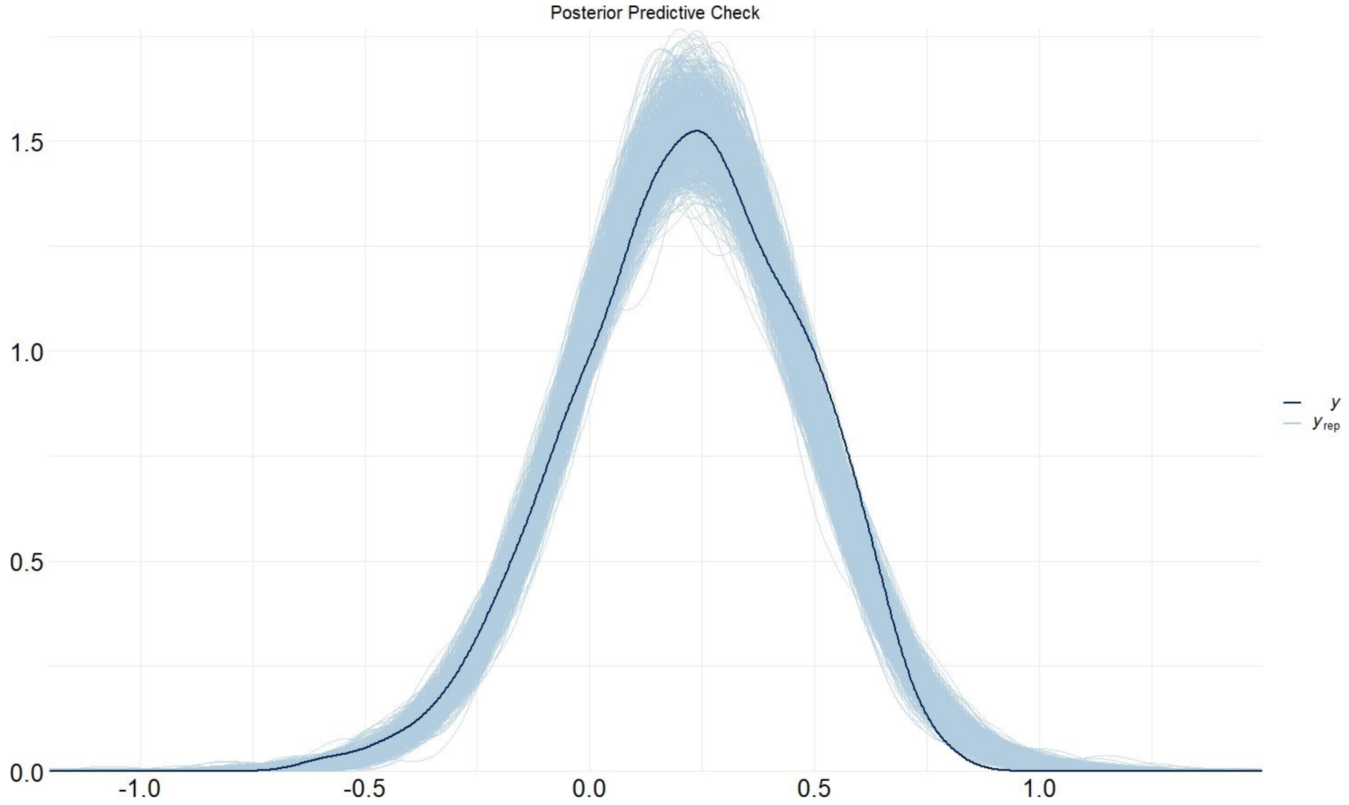


Supplementary Figure 8 : Posterior predictive draws (cyan) of 1000 simulations from the fitted model compared with data (black) for the information encapsulation Bayesian model for intersubject correlation (ISC) analysis within prefrontal regions during domain updates. The model had (Bayes adjusted) $R^2$ of 0.48.

### Supplementary Fig 9

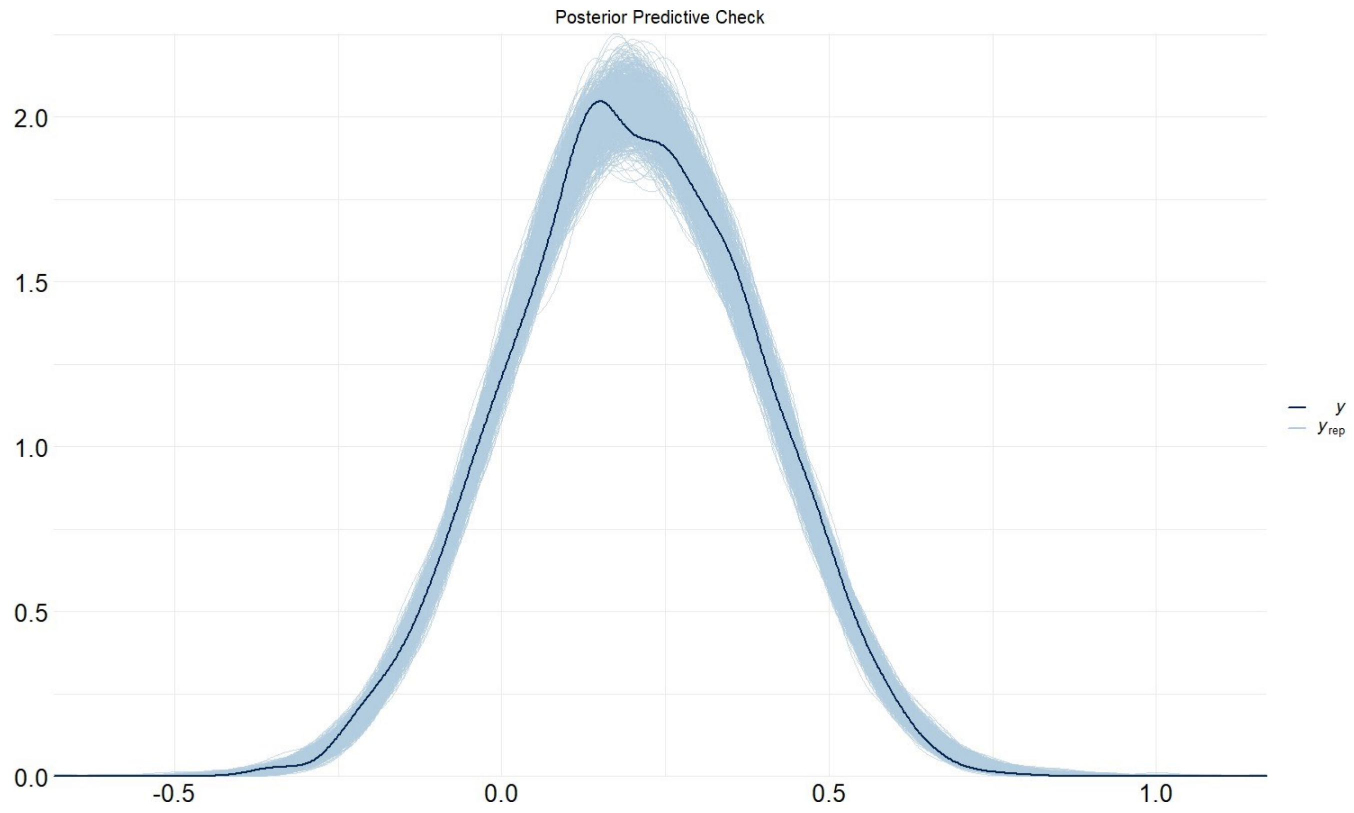


Supplementary Figure 9 : Posterior predictive draws (cyan) of 1000 simulations from the fitted model compared with data (black) for the integration Bayesian model for intersubject functional correlation (ISFC) analysis between regions during domain updates. The model had (Bayes adjusted) $R^2$ of 0.85.

### Supplementary Fig 10

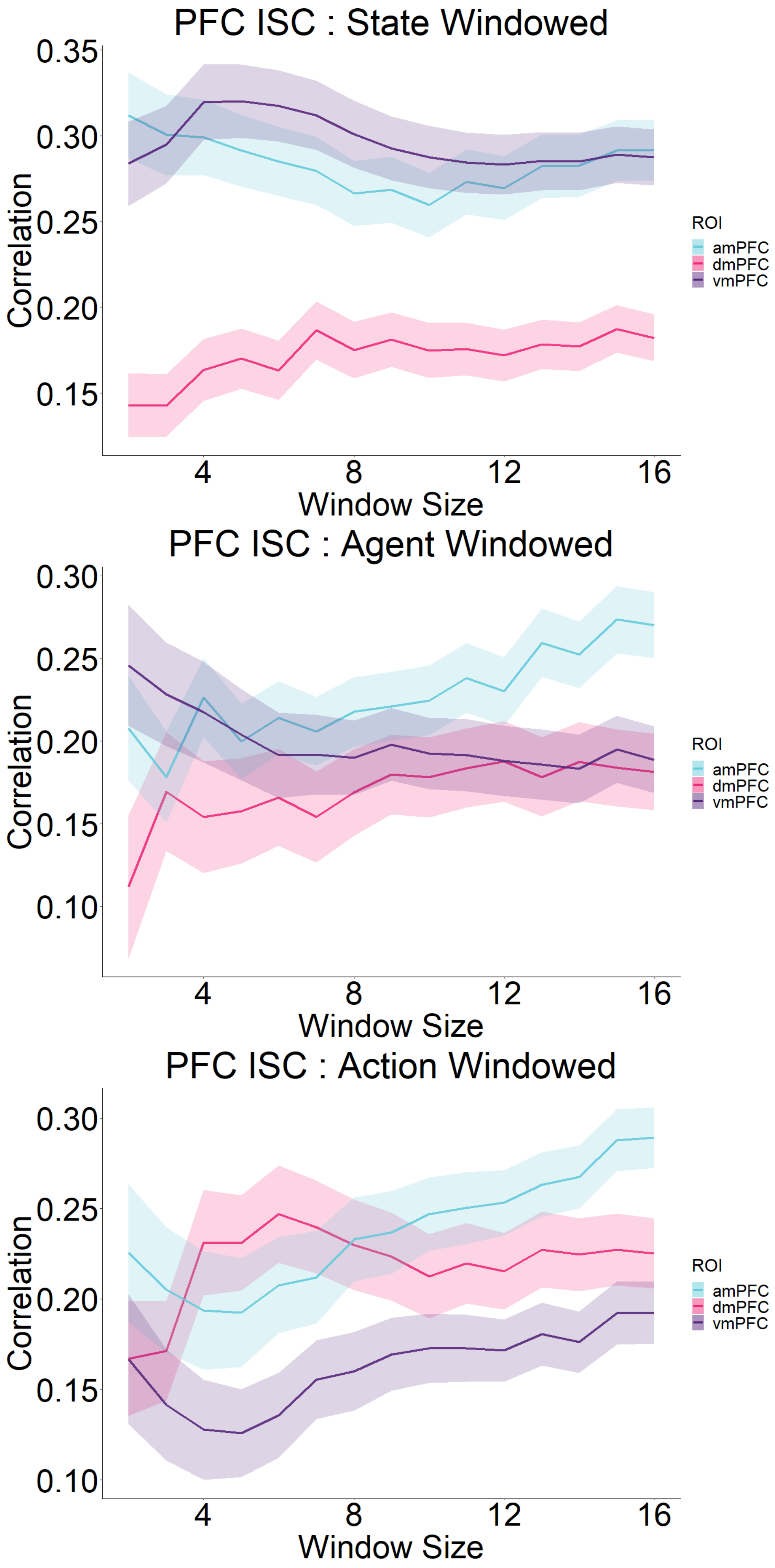
PFC ISC : State Windowed
0.35
0.30
0.25
0.20
0.15
Correlation
4
8
12
16
Window Size
ROI
amPFC
dmPFC
vmPFC
PFC ISC : Agent Windowed
0.30
0.25
0.20
0.15
0.10
Correlation
4
8
12
16
Window Size
ROI
amPFC
dmPFC
vmPFC
PFC ISC : Action Windowed
0.30
0.25
0.20
0.15
0.10
Correlation
4
8
12
16
Window Size
ROI
amPFC
dmPFC
vmPFC

Supplementary  Figure 10 : Window analysis ISC. ISC analysis in Fig 3c in the main text repeated across a range of (around) windows. Main text used 7TR window. Bands show standard error of the mean.

#### Supplementary Fig 11

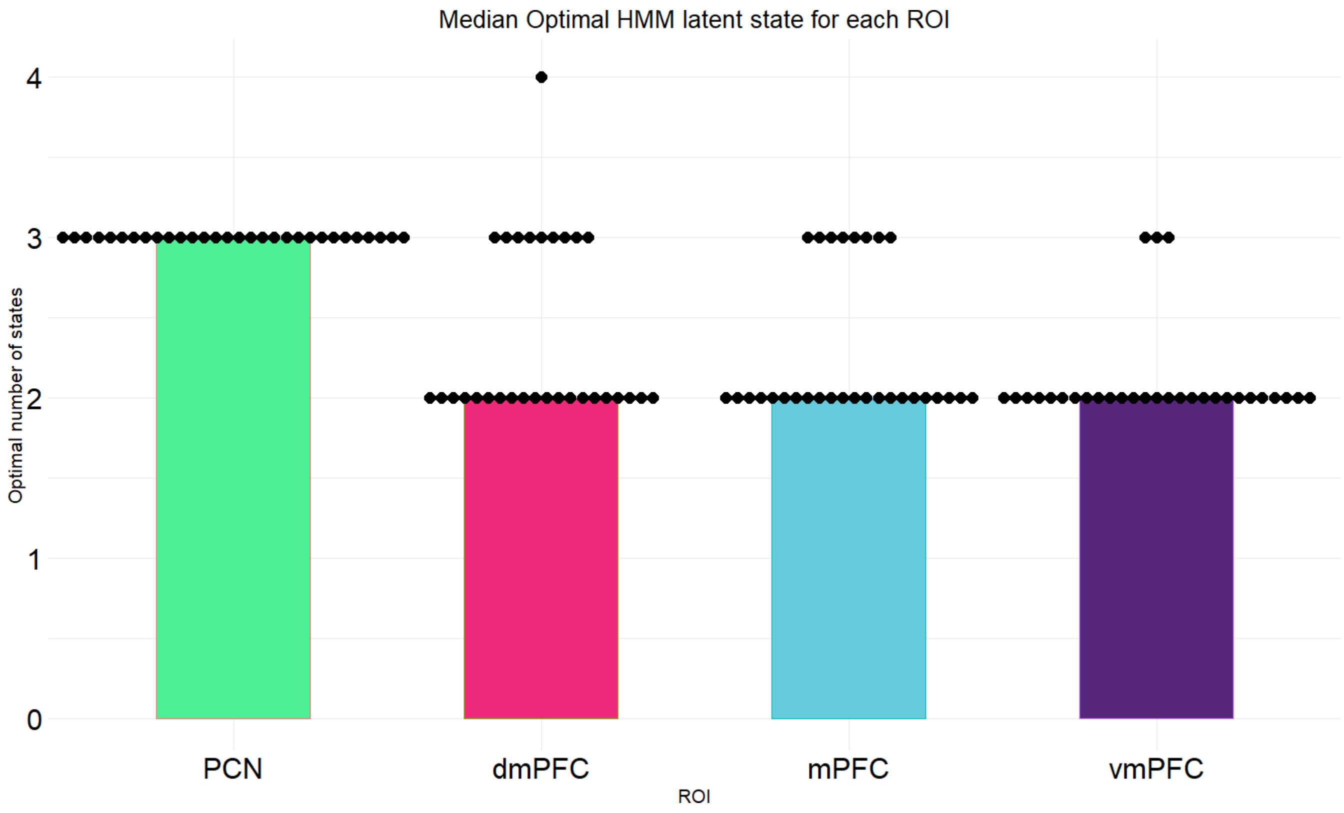


Supplementary  Figure 11 : Optimal number of latent states for each ROI during each of the 30 runs (dots) and bars represent the median latent state in each.

### Supplementary Fig 12

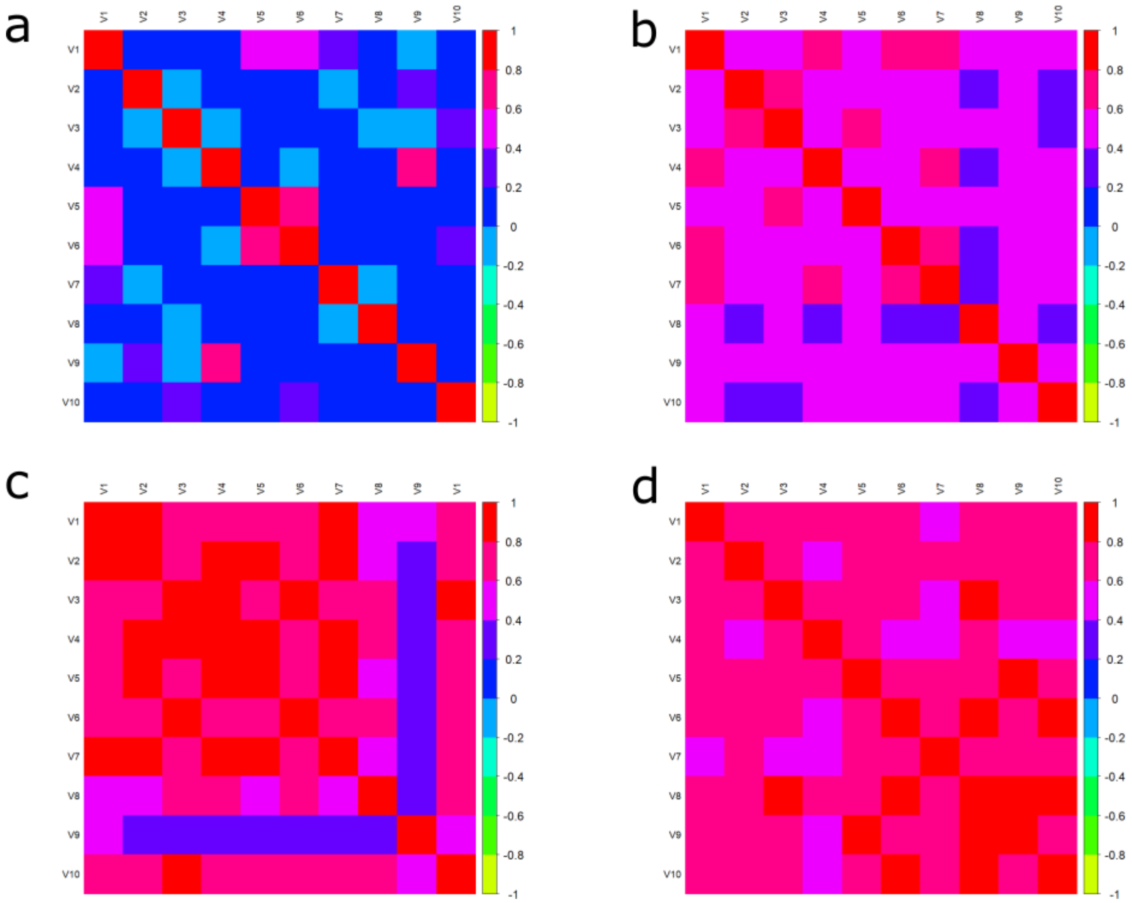


Supplementary  Figure 12 : Inter-run correlation of the transition time-series on the final 10 runs of HMM in a) dmPFC b) amPFC c) Precuneus and d) vmPFC.

### Supplementary Fig 13

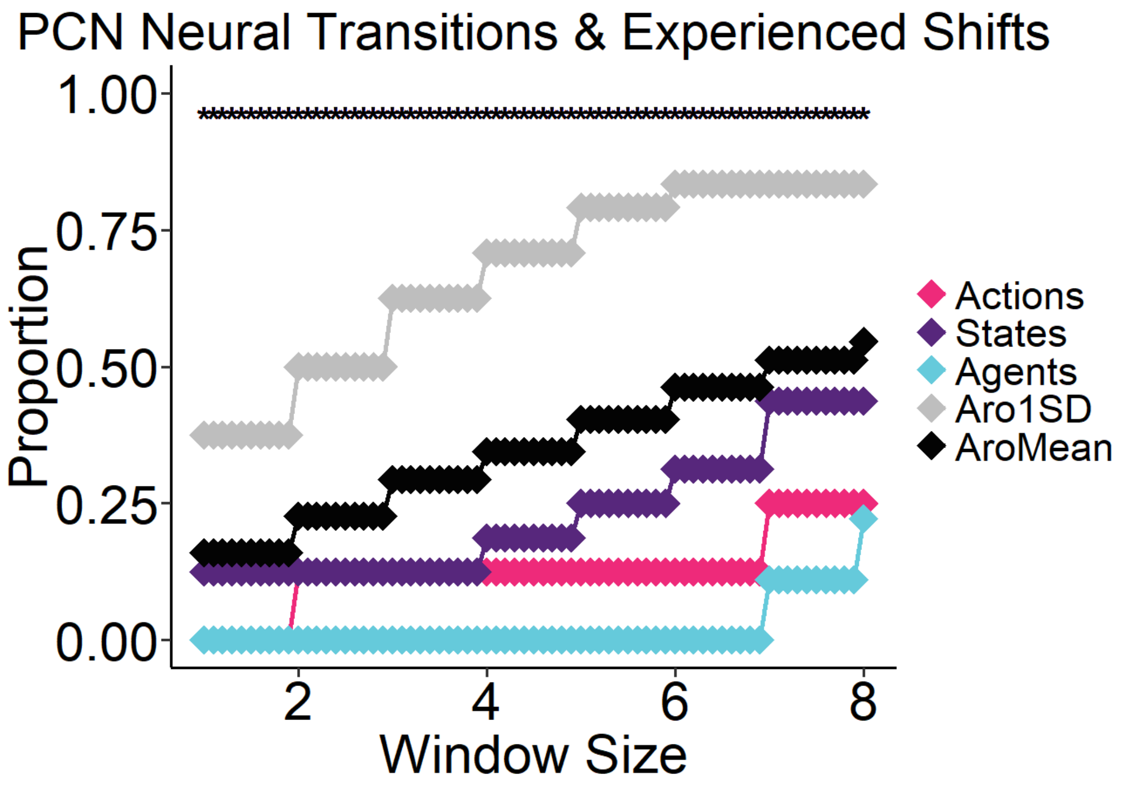


Supplementary  Figure 13 : HMM neural shift analysis of Precuneus with Arousal shifts with  θ set to mean (black) compared to 1SD (grey) which was used in the main analysis

#### Supplementary Fig 14

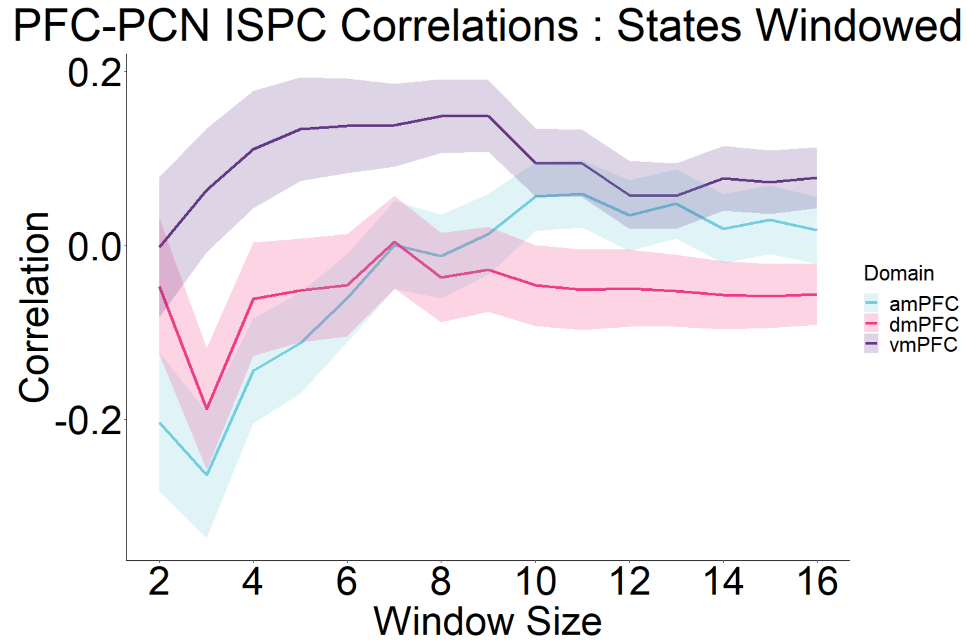
PFC-PCN ISPC Correlations : States Windowed
0.2
0.0
-0.2
Correlation
2
4
6
8
10
12
14
16
Window Size
Domain
amPFC
dmPFC
vmPFC

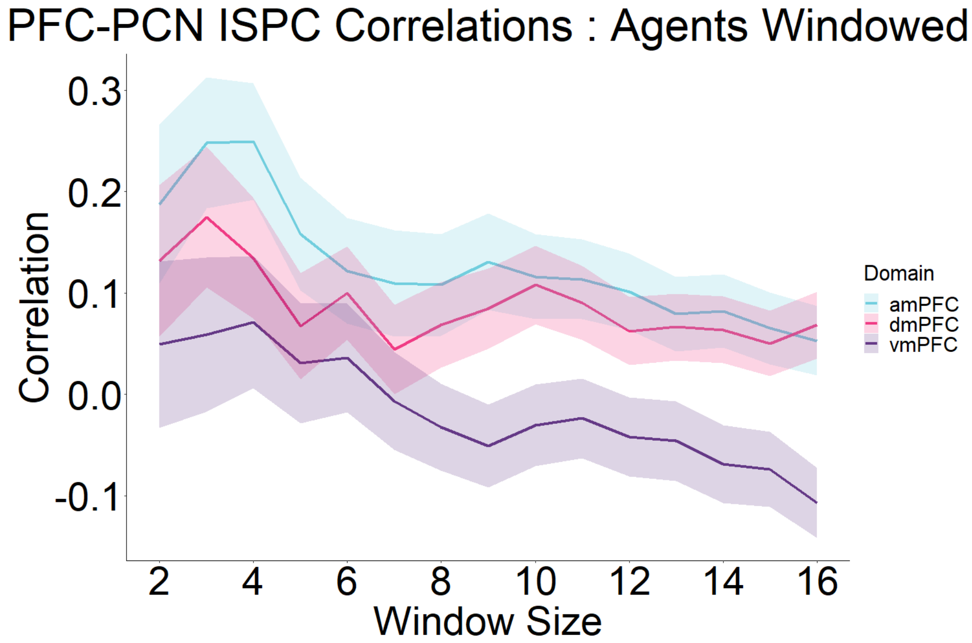
PFC-PCN ISPC Correlations : Agents Windowed
0.3
0.2
0.1
0.0
-0.1
Correlation
2
4
6
8
10
12
14
16
Window Size
Domain
amPFC
dmPFC
vmPFC

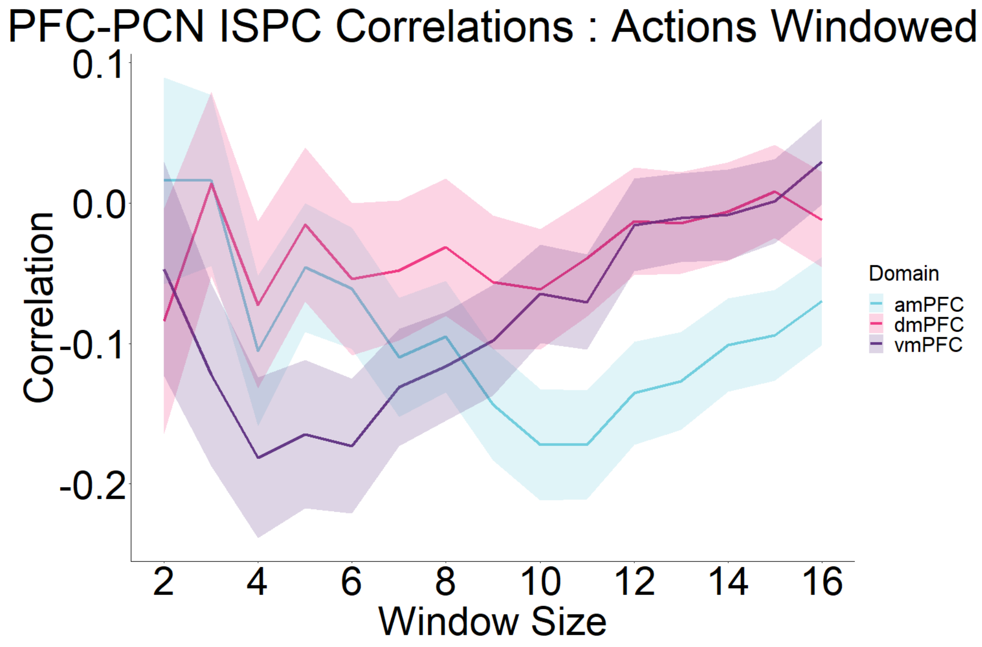
PFC-PCN ISPC Correlations : Actions Windowed
0.1
0.0
-0.1
-0.2
Correlation
2
4
6
8
10
12
14
16
Window Size
Domain
amPFC
dmPFC
vmPFC

Supplementary Figure 14 : Window analysis ISPC. ISPC analysis in Fig 5d in the main text repeated across a range of (around) windows. Main text used 9TR window. Bands show standard error of the mean.

#### Supplementary Fig 15

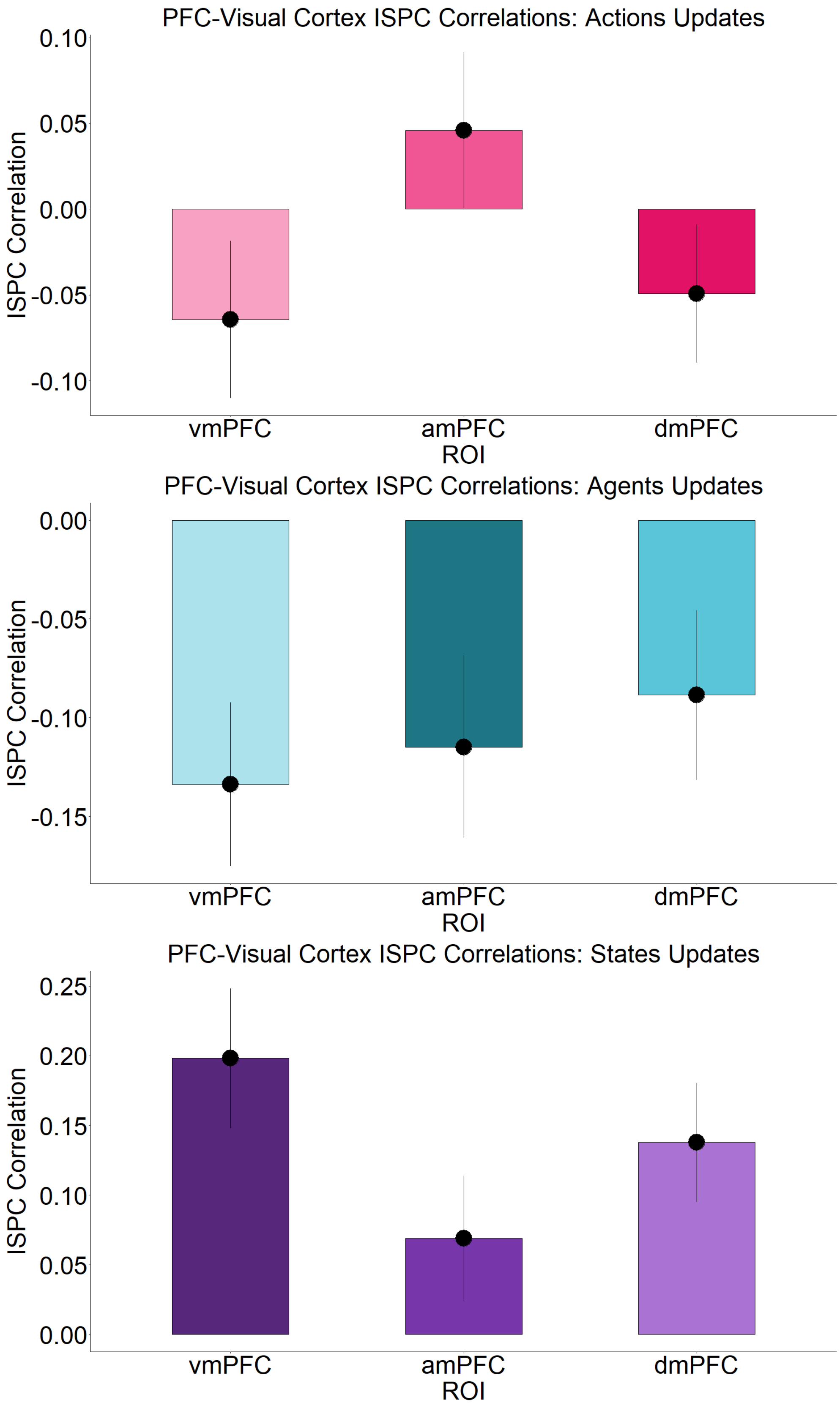

PFC-Visual Cortex ISPC Correlations: Actions Updates
ISPC Correlation
0.10
0.05
0.00
-0.05
-0.10
vmPFC
amPFC
dmPFC
ROI
PFC-Visual Cortex ISPC Correlations: Agents Updates
ISPC Correlation
0.00
-0.05
-0.10
-0.15
vmPFC
amPFC
dmPFC
ROI
PFC-Visual Cortex ISPC Correlations: States Updates
ISPC Correlation
0.25
0.20
0.15
0.10
0.05
0.00
vmPFC
amPFC
dmPFC
ROI

Supplementary Figure 15: Multithreading analysis in Visual Cortex. Bar plots show correlation strength of region's ISPC timecourse during updates with that of each PFC during updates. Error bars show standard error of the mean.

#### Supplementary Fig 16

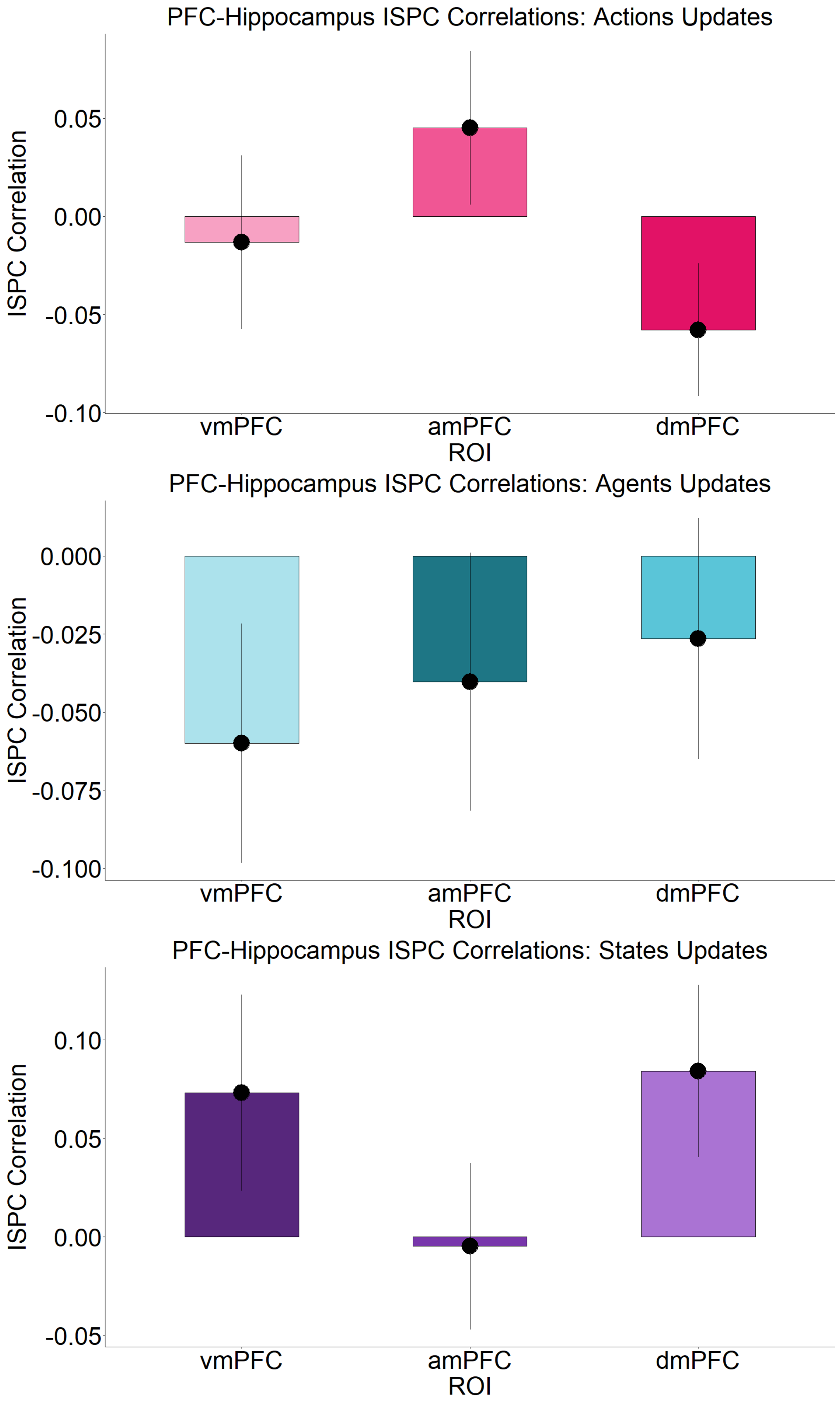
PFC-Hippocampus ISPC Correlations: Actions Updates
ISPC Correlation
0.05
0.00
-0.05
-0.10
vmPFC
amPFC
dmPFC
ROI
PFC-Hippocampus ISPC Correlations: Agents Updates
ISPC Correlation
0.000
-0.025
-0.050
-0.075
-0.100
vmPFC
amPFC
dmPFC
ROI
PFC-Hippocampus ISPC Correlations: States Updates
ISPC Correlation
0.10
0.05
0.00
-0.05
vmPFC
amPFC
dmPFC
ROI

Supplementary Figure 16 : Multithreading analysis in Hippocampus. Bar plots show correlation strength of region's ISPC timecourse during updates with that of each PFC during updates. Error bars show standard error of the mean.

#### Supplementary Fig 17

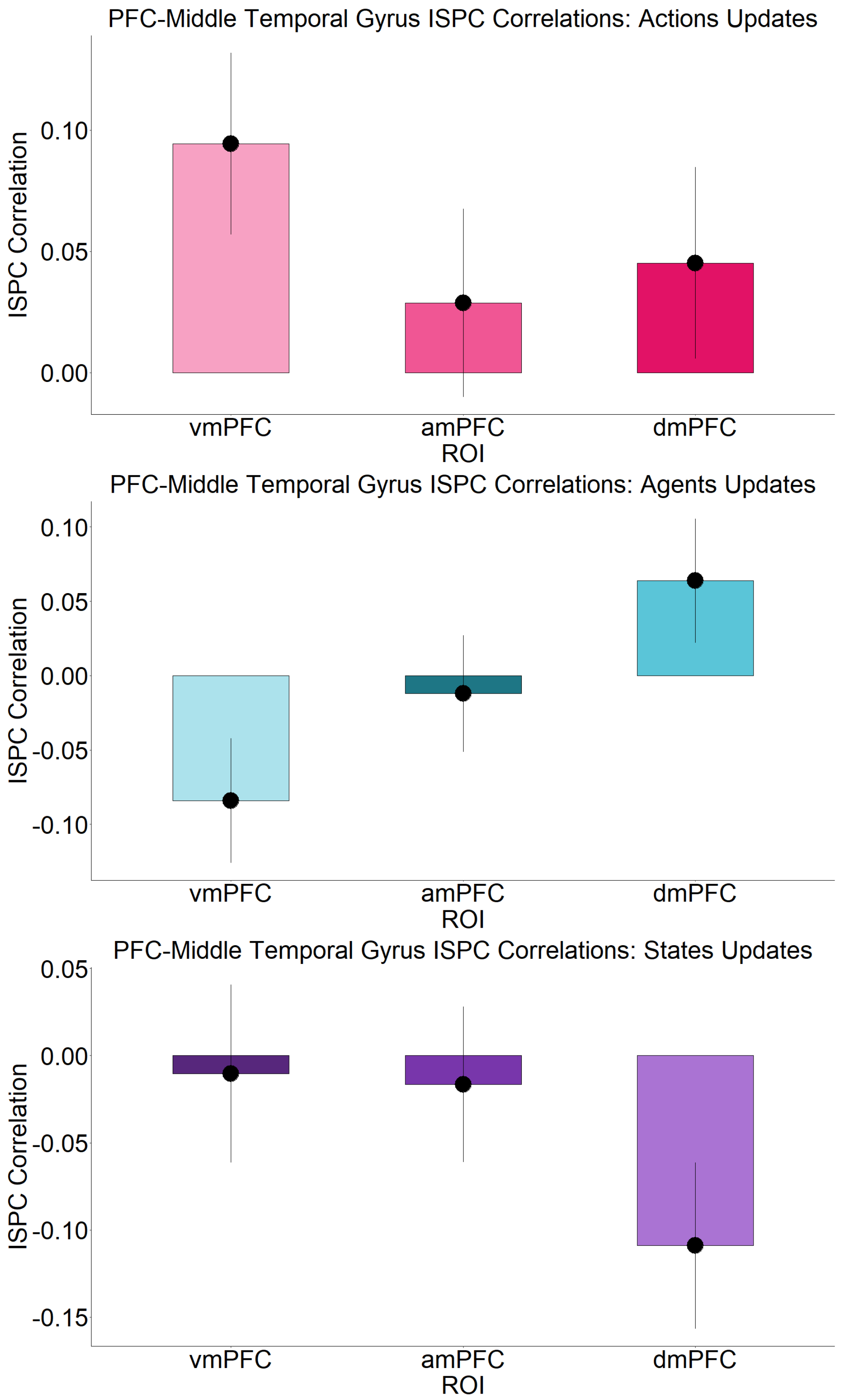
PFC-Middle Temporal Gyrus ISPC Correlations: Actions Updates
ISPC Correlation
0.10
0.05
0.00
vmPFC
amPFC
dmPFC
ROI
PFC-Middle Temporal Gyrus ISPC Correlations: Agents Updates
ISPC Correlation
0.10
0.05
0.00
-0.05
-0.10
vmPFC
amPFC
dmPFC
ROI
PFC-Middle Temporal Gyrus ISPC Correlations: States Updates
ISPC Correlation
0.05
0.00
-0.05
-0.10
-0.15
vmPFC
amPFC
dmPFC
ROI

Supplementary Figure 17 : Multithreading analysis in Middle Temporal Gyrus. Bar plots show correlation strength of region's ISPC timecourse during updates with that of each PFC during updates. Error bars show standard error of the mean.

#### Supplementary Fig 18

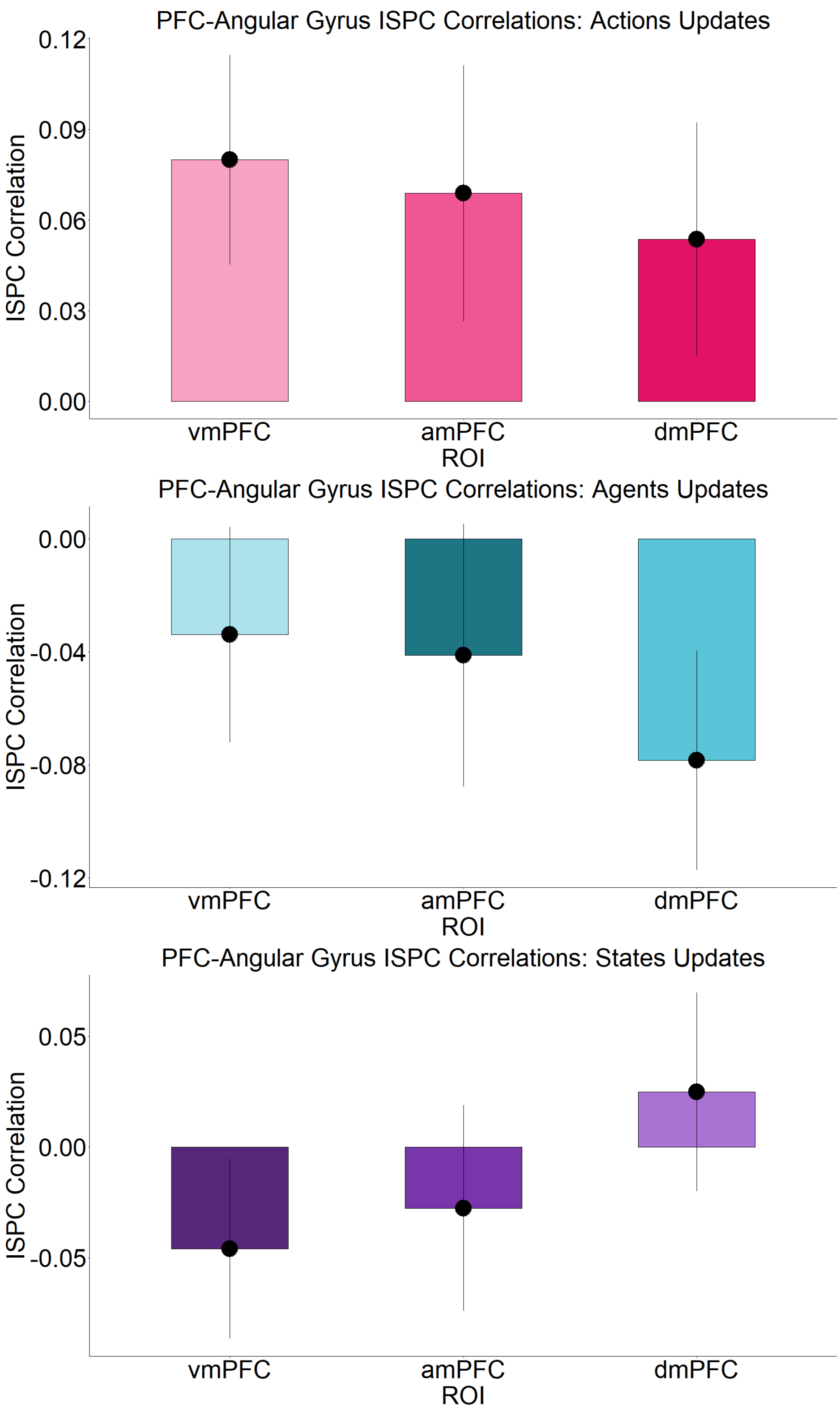
PFC-Angular Gyrus ISPC Correlations: Actions Updates
ISPC Correlation
0.12
0.09
0.06
0.03
0.00
vmPFC
amPFC
dmPFC
ROI
PFC-Angular Gyrus ISPC Correlations: Agents Updates
ISPC Correlation
0.00
-0.04
-0.08
-0.12
vmPFC
amPFC
dmPFC
ROI
PFC-Angular Gyrus ISPC Correlations: States Updates
ISPC Correlation
0.05
0.00
-0.05
vmPFC
amPFC
dmPFC
ROI

Supplementary Figure 18 : Multithreading analysis in Angular Gyrus. Bar plots show correlation strength of region's ISPC timecourse during updates with that of each PFC during updates. Error bars show standard error of the mean.

#### Supplementary Fig 19

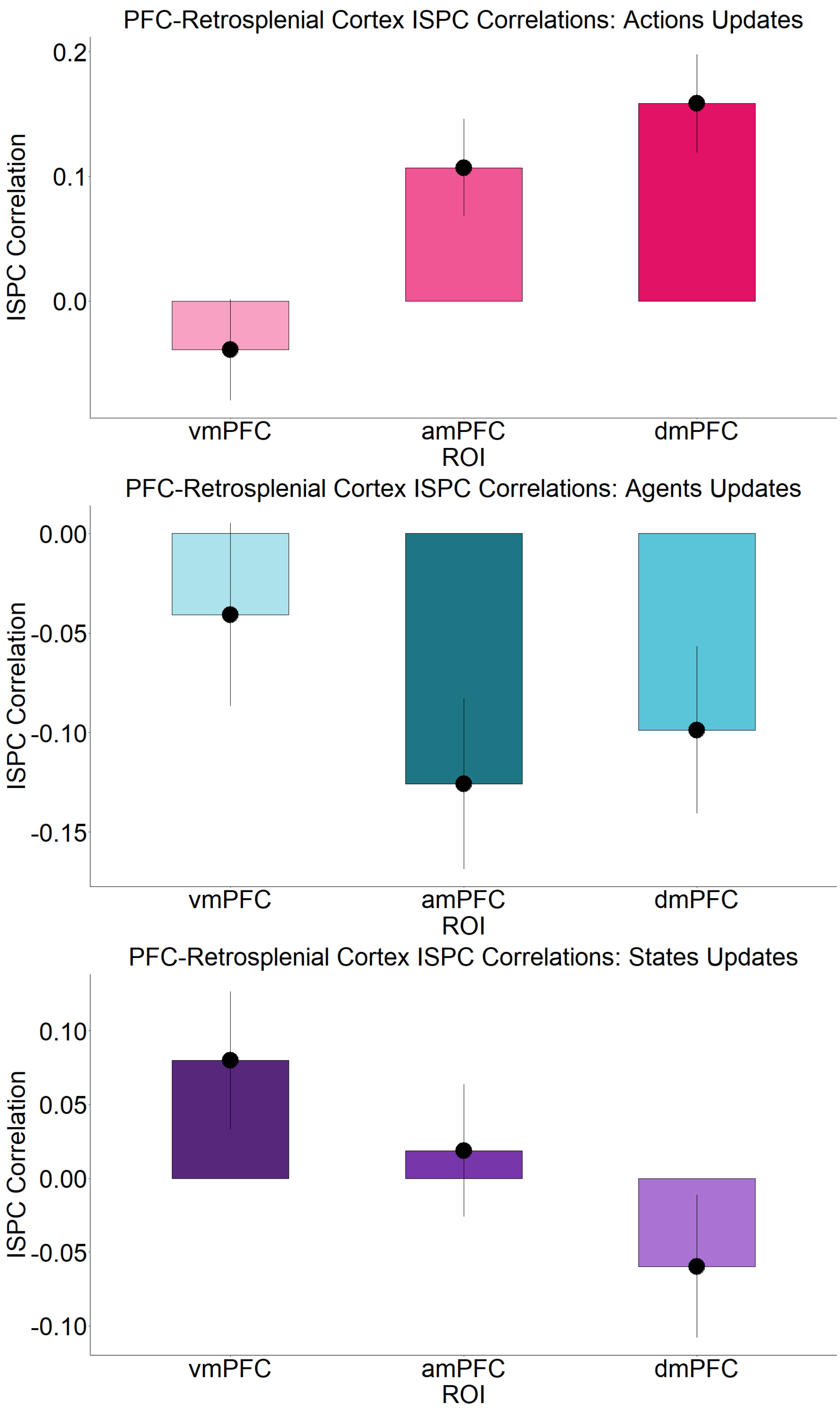
PFC-Retrosplenial Cortex ISPC Correlations: Actions Updates
ISPC Correlation
0.2
0.1
0.0
vmPFC
amPFC
dmPFC
ROI
PFC-Retrosplenial Cortex ISPC Correlations: Agents Updates
ISPC Correlation
0.00
-0.05
-0.10
-0.15
vmPFC
amPFC
dmPFC
ROI
PFC-Retrosplenial Cortex ISPC Correlations: States Updates
ISPC Correlation
0.10
0.05
0.00
-0.05
-0.10
vmPFC
amPFC
dmPFC
ROI

Supplementary Figure 19 : Multithreading analysis in Retrosplenial Cortex. Bar plots show correlation strength of region's ISPC timecourse during updates with that of each PFC during updates. Error bars show standard error of the mean.

#### Supplementary Fig 20

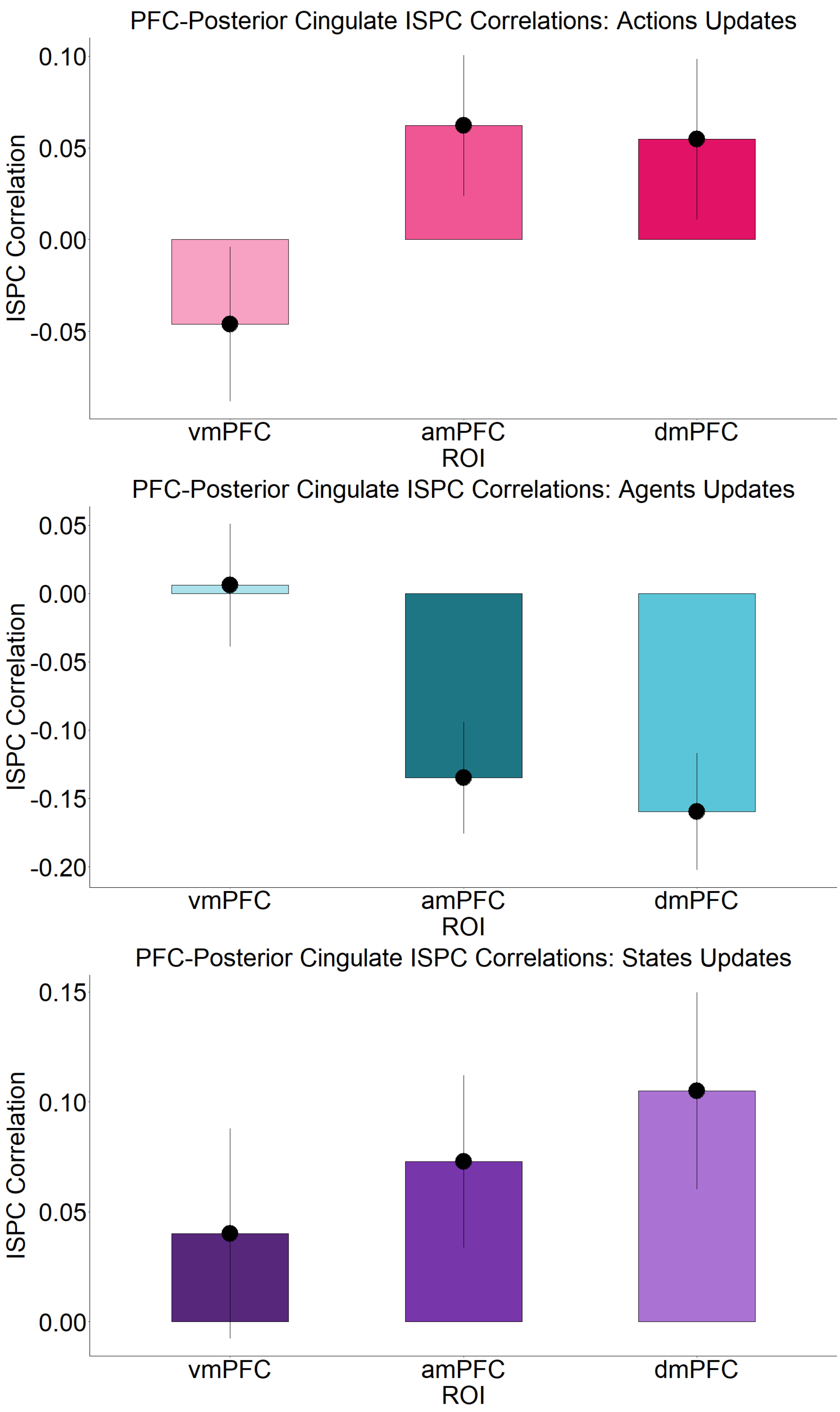
PFC-Posterior Cingulate ISPC Correlations: Actions Updates
ISPC Correlation
0.10
0.05
0.00
-0.05
vmPFC
amPFC
dmPFC
ROI
PFC-Posterior Cingulate ISPC Correlations: Agents Updates
ISPC Correlation
0.05
0.00
-0.05
-0.10
-0.15
-0.20
vmPFC
amPFC
dmPFC
ROI
PFC-Posterior Cingulate ISPC Correlations: States Updates
ISPC Correlation
0.15
0.10
0.05
0.00
vmPFC
amPFC
dmPFC
ROI

Supplementary Figure 20 : Multithreading analysis in Posterior Cingulate Cortex. Bar plots show correlation strength of region's ISPC timecourse during updates with that of each PFC during updates. Error bars show standard error of the mean.

#### Supplementary Fig 21

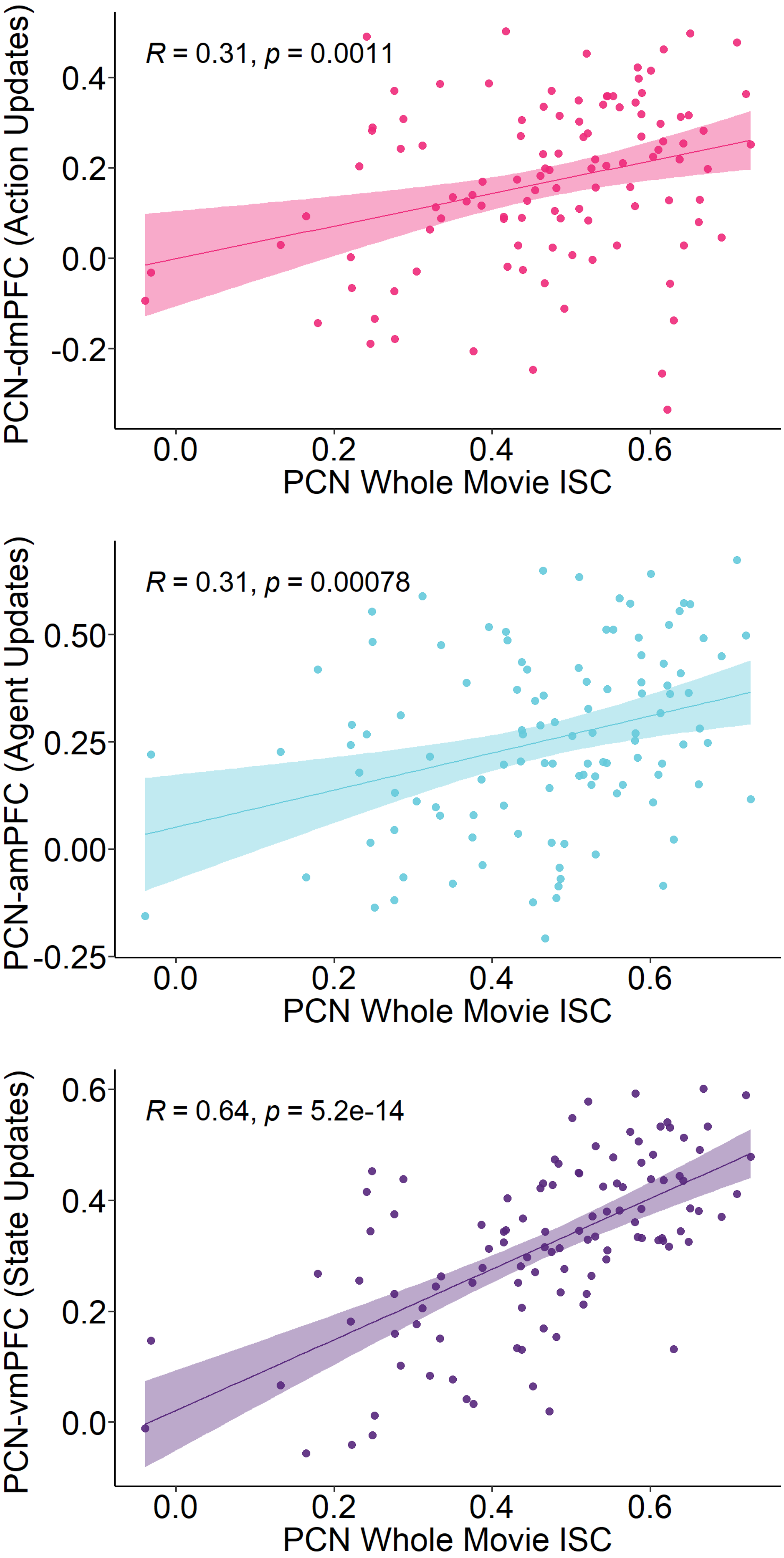
R = 0.31, p = 0.0011
PCN-dmPFC (Action Updates)
0.4
0.2
0.0
-0.2
0.0
0.2
0.4
0.6
PCN Whole Movie ISC
R = 0.31, p = 0.00078
PCN-amPFC (Agent Updates)
0.50
0.25
0.00
-0.25
0.0
0.2
0.4
0.6
PCN Whole Movie ISC
R = 0.64, p = 5.2e-14
PCN-vmPFC (State Updates)
0.6
0.4
0.2
0.0
0.0
0.2
0.4
0.6
PCN Whole Movie ISC

Supplementary Figure 21 : Scatterplots show average functional connectivity between Precuneus and (top) dmPFC,(middle) amPFC and vmPFC, during updates (to Actions, Agents States respectively) along y-axis, correlated with the whole-movie Precuneus ISC on x-axis. Dots represent individual participants.

### Supplementary Fig 22

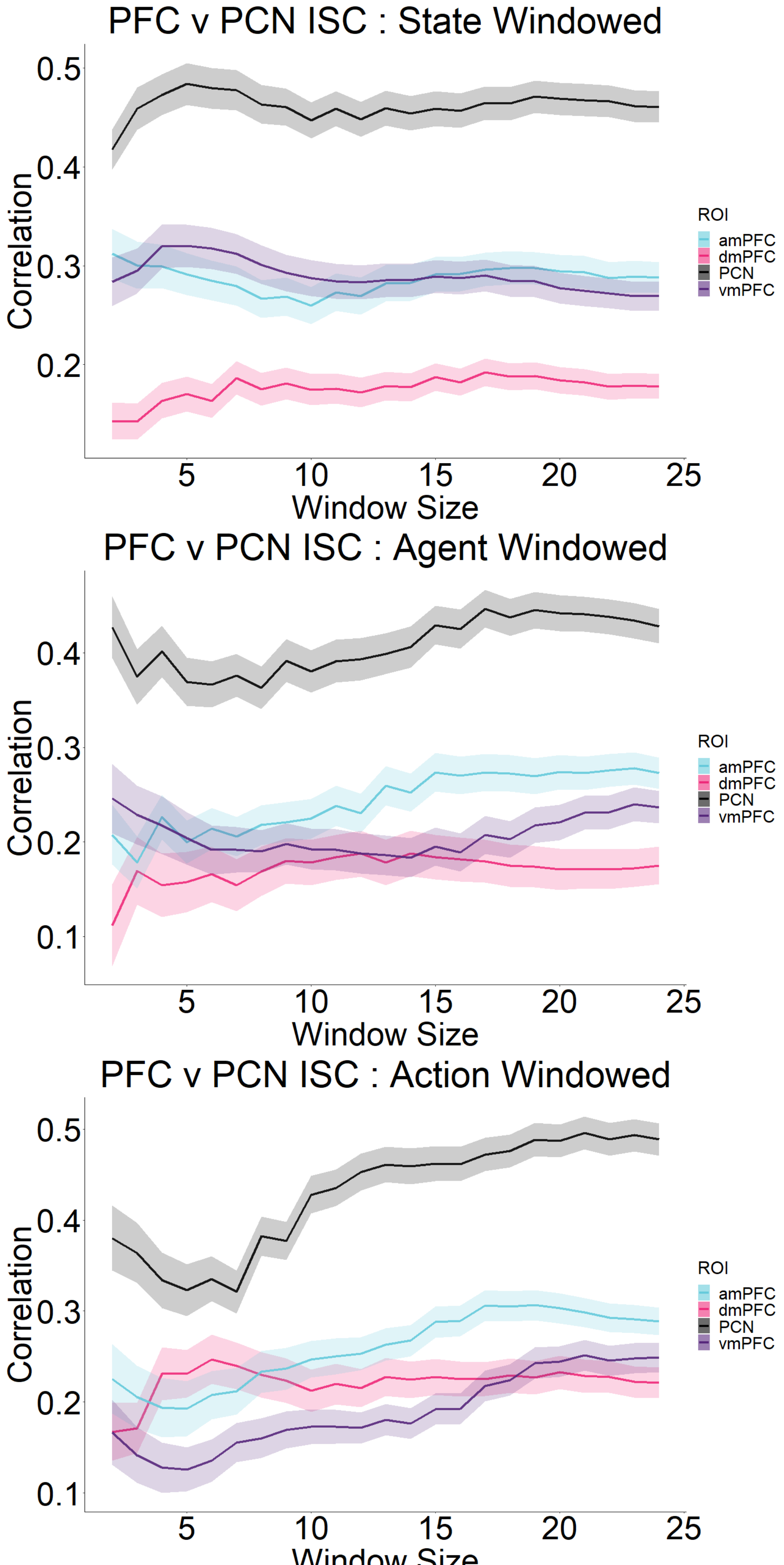
PFC v PCN ISC : State Windowed
Correlation
0.5
0.4
0.3
0.2
5
10
15
20
25
Window Size
ROI
amPFC
dmPFC
PCN
vmPFC
PFC v PCN ISC : Agent Windowed
Correlation
0.4
0.3
0.2
0.1
5
10
15
20
25
Window Size
ROI
amPFC
dmPFC
PCN
vmPFC
PFC v PCN ISC : Action Windowed
Correlation
0.5
0.4
0.3
0.2
0.1
5
10
15
20
25
ROI
amPFC
dmPFC
PCN
vmPFC

Supplementary Figure 22 : Window analysis ISC over longer windows with PFCs and Precuneus. This is the same as Supplementary Figure 10, but the (around) windows now extend to 1 min (25 TR or ~60s). Bands show standard error of the mean.

### Supplementary Fig 23

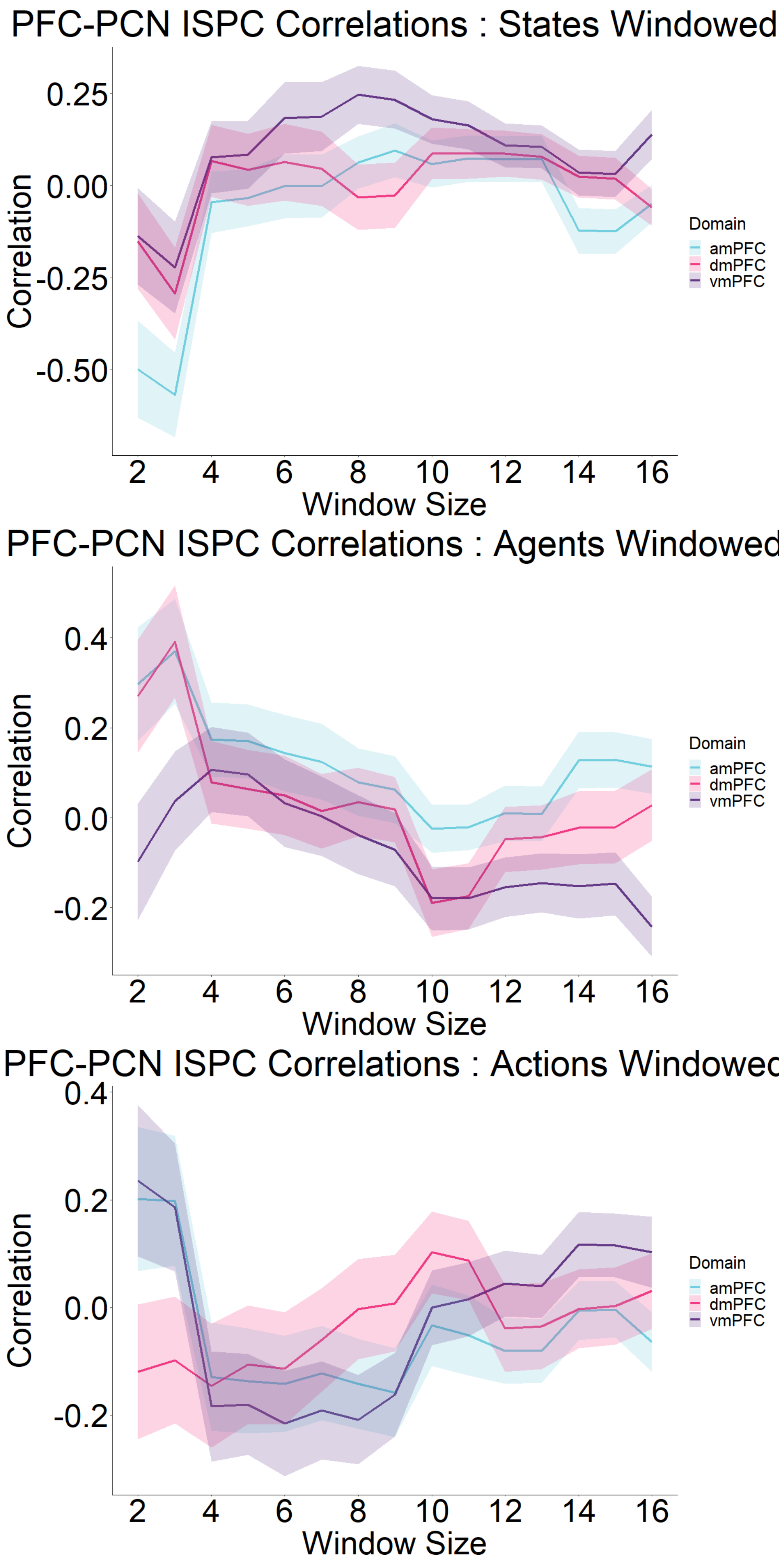
PFC-PCN ISPC Correlations : States Windowed
Correlation
0.25
0.00
-0.25
-0.50
2
4
6
8
10
12
14
16
Window Size
Domain
amPFC
dmPFC
vmPFC
PFC-PCN ISPC Correlations : Agents Windowed
Correlation
0.4
0.2
0.0
-0.2
2
4
6
8
10
12
14
16
Window Size
Domain
amPFC
dmPFC
vmPFC
PFC-PCN ISPC Correlations : Actions Windowec
Correlation
0.4
0.2
0.0
-0.2
2
4
6
8
10
12
14
16
Window Size
Domain
amPFC
dmPFC
vmPFC

Supplementary  Figure 23 : Window analysis ISPC in Narrative. ISPC analysis in Fig 5d in the main text repeated across a range of (around) windows. Main text used 9TR window. Bands show standard error of the mean.

### Supplementary Fig 24

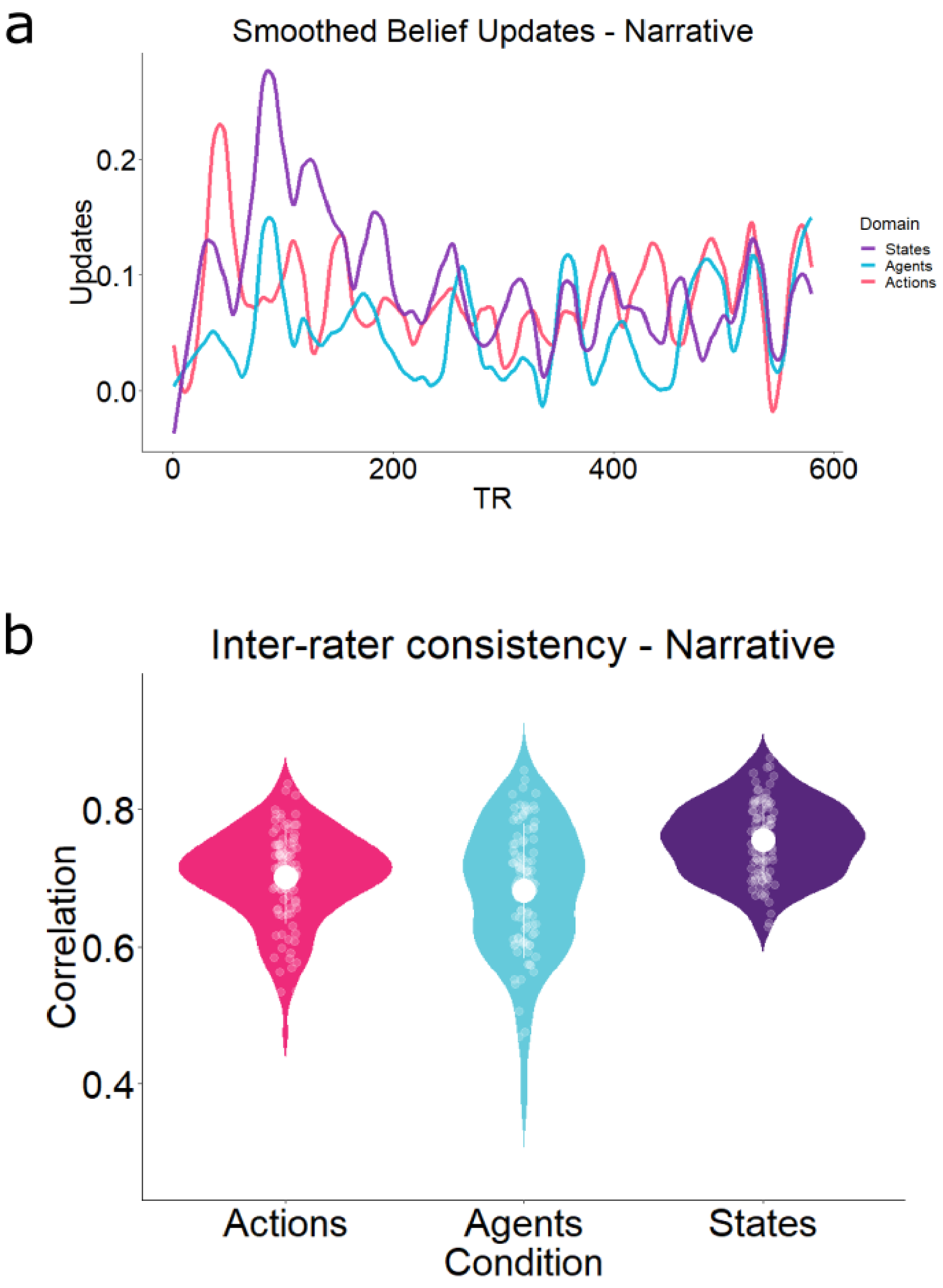
a
Smoothed Belief Updates - Narrative
Updates
0.2
0.1
0.0
0
200
400
600
TR
Domain
States
Agents
Actions
b
Inter-rater consistency - Narrative
Correlation
0.8
0.6
0.4
Actions
Agents
States
Condition

Supplementary  Figure 24 : a) Smoothed prediction updates for the Narrative and b) their inter-rater reliability analysis conducted by split half correlation, adjusted through Spearman Brown formula. States had r = 0.75, Agents r = 0.68 and Actions r = 0.70.
